# Cyclic pathogen epidemics favour the evolution of delayed germination

**DOI:** 10.64898/2026.09.04.749465

**Authors:** Mahdi Salehzadeh, John M. Stockie, Ailene MacPherson

## Abstract

Delayed germination is classically explained as a bet-hedging strategy against environmental variability: by withholding a fraction of seeds from germination, plants spread establishment risk across years that vary in temperature, precipitation, and other conditions governing seedling survival. Whether biotic interactions can generate analogous selective pressure has received comparatively little theoretical attention. Here we examine the evolution of germination timing in a host-pathogen model in which a saturating (Holling Type II) transmission function couples seed bank dynamics to epidemic cycles. The model produces three qualitatively distinct ecological regimes: a pathogen-free equilibrium, a stable endemic equilibrium, and an oscillatory endemic regime arising through a bifurcation, a transition from stable to oscillatory endemic dynamics. Using adaptive dynamics, namely invasion analysis, we show that the evolutionarily stable germination rate depends critically on which regime the resident population occupies. When pathogen dynamics settle to an equilibrium, selection favours ever-faster germination with no finite optimum, regardless of pathogen presence. When dynamics are limit cycles, periodic epidemic peaks create recurrent windows of high establishment mortality that function as biotic analogues of abiotic interactions (environmental), and selection drives the germination rate toward the bifurcation boundary. The evolutionary attractor thus coincides with an ecological bifurcation point. Trait substitution sequence simulations confirm convergence to this attractor, and multi-trait eco-evolutionary simulations provide evidence that the attractor is evolutionarily stable, consistent with a continuously stable strategy (CSS). These results extend classical bet-hedging theory to biotic drivers and suggest that pathogens capable of sustaining population cycles may be an underappreciated selective force on germination timing and, more broadly, on the pace of life-history evolution.

## 1 Introduction

Many organisms paradoxically delay reproduction even though, all else being equal, earlier reproduction increases fitness (Tuljapurkar and Wiener, 2000). Seed dormancy in annual plants is a canonical example (Cohen, 1966). Early theoretical work established that environmental stochasticity resolves the paradox: when abiotic conditions fluctuate between favourable and unfavourable seasons, a seed that does not germinate immediately but retains the option to do so in a future season, thereby spreading the risk of establishment failure across time rather than committing all reproductive effort to a single, potentially poor year (Scott and Otto, 2014). This temporal risk-spreading has a precise population-genetic interpretation: because natural selection in variable environments acts on the geometric mean fitness rather than the arithmetic mean, delayed germination can be selectively maintained even when it reduces expected reproductive output within any single generation (Scott and Otto, 2014). The existing literature largely attributes environmental variability to abiotic sources. Cyclic pathogen prevalence generates recurrent seasons of high infection risk that parallel the role of abiotic bad years in classical bet-hedging models. Whether such biotic forcing selects for, or against, delayed germination, and at what germination rate an evolutionarily stable strategy emerges, form the central questions of the present study.

The formal theory of delayed seed germination was set out by Cohen (1966), who showed that in an environment where years vary independently in their suitability for germination, the evolutionarily optimal germination fraction is governed by the ratio of dormant-seed survival to the probability that any given year is favourable. When this ratio exceeds one, partial dormancy is maintained by selection; when it falls below one, immediate germination is optimal. Subsequent theoretical work extended this framework to incorporate spatial heterogeneity and dispersal (Venable and Brown, 1988), age-structured seed banks (Rees, 1994), and finer decompositions of environmental variance (Gremer and Venable, 2014). Across this body of theory, the source of variability driving dormancy evolution has consistently been treated as exogenous: precipitation regimes, temperature fluctuations, or disturbance events that originate outside the focal population (Finch-Savage and LeubnerMetzger, 2006).

This abiotic emphasis is well-justified in many natural systems, where year-to-year climatic variation is the dominant driver of establishment success (Gremer and Venable, 2014). It nevertheless leaves an important class of selective agent unexplored. The biotic interactions, including competitors, herbivores, and pathogens, is itself a dynamical quantity that fluctuates in time, and if that fluctuation is sufficiently structured and large in amplitude it could, in principle, generate selection on germination timing analogous to that produced by abiotic stochasticity.

Among the biotic interactions capable of producing temporally structured establishment risk, pathogens occupy a prominent position. Rather than acting as a constant background hazard, pathogen may effectively create recurrently unfavourable periods for reproduction: cohorts of seeds that germinate during phases of high infection prevalence suffer elevated mortality, whereas those that germinate during low-prevalence phases enjoy greater establishment success. Periodicity in pathogen prevalence can arise through a variety of mechanisms, and here we focus on one: density-dependent transmission rates. Specifically, when the per-capita transmission rate saturates at high host density, as described by a Holling Type II functional response (Holling, 1959), the coupling between host abundance and infection pressure becomes nonlinear. This saturating transmission mechanism is therefore capable of generating sustained, cyclic fluctuations in pathogen prevalence even in the absence of any external environmental forcing. These endogenously generated cycles of high and low infection risk are structurally analogous to the exogenous abiotic fluctuations that classical theory identifies as the selective driver of delayed seed germination, and it is this analogy that motivates the present study.

The cyclic pathogen dynamics described above arise naturally from the nonlinearity of the infection process. At low host density, pathogen spread is contact-limited and the percapita infection rate rises approximately linearly with host abundance. At high host density, however, each pathogen unit is occupied by processing current infections for a finite handling time, so the effective transmission rate saturates, a phenomenon analogous to the Holling Type II functional response in predator-prey theory (Holling, 1959). In the epidemiological context, handling time captures the duration of the infectious period or the time required for a pathogen to complete its life cycle within a single host. Mathematically speaking, incorporating this saturation into a susceptible-infected host model with logistic host growth produces a system capable of limit cycles in infected host density. The occurrence and amplitude of the epidemic cycles depend on the transmission rate, the host carrying capacity, and the handling time. We expect magnitude and frequency of the resulting epidemic cycles to modulate the sign and strength of selection acting on the germination rate. As in Scott and Otto (2014) we use an adaptive dynamics/invasion analysis approach to explore the resulting evolution of germination rate.

From an adaptive dynamics perspective, what makes this system particularly interesting is that the germination rate, *g*, the trait under selection, is also the bifurcation parameter governing the resident ecological dynamics (i.e., the parameter whose value determines the qualitative character of the long-run dynamics). Depending on where *g* sits relative to the Hopf bifurcation point (the threshold value of *g* at which the endemic equilibrium loses stability and gives way to limit cycles), the resident population either approaches a stable endemic equilibrium or settles into a sustained limit cycle of infection prevalence. As the trait evolves, the nature of the environment experienced by rare mutants changes accordingly. When the resident population dynamics are characterized by a limit cycle, classical invasion analysis, which evaluates mutant growth against a fixed equilibrium, is no longer applicable. Invasion fitness must instead be computed by tracking a rare mutant cohort through one complete oscillation of the resident-driven cycle, and Floquet theory provides the natural framework for doing so: the dominant Floquet multiplier of the linearised mutant subsystem over one period determines whether the mutant can increase when rare (Klausmeier, 2008; Best and Ashby, 2023). Embedding this fitness criterion within the adaptive dynamics framework then allows us to identify convergence-stable, uninvadable germination rates as the long-run evolutionary outcome, and to ask how that outcome shifts as ecological parameters move the resident dynamics across the bifurcation.

This paper proceeds by first establishing the ecological foundation: we introduce a host-pathogen model with a seed bank and characterise how the germination rate governs the qualitative nature of the resident dynamics, producing three distinct regimes separated by a Hopf bifurcation. We then embed this ecological structure within an evolutionary framework, deriving the invasion fitness of a rare mutant through Floquet analysis and mapping the selective landscape across the ecological parameter space. Adaptive dynamics then identifies the evolutionarily stable germination rate, and the central result is that this evolutionary attractor coincides with the Hopf bifurcation boundary; that is, selection drives the population to the edge of cyclic dynamics, where pathogen prevalence oscillates but does not explode. Numerical trait substitution sequence simulations confirm convergence to this attractor from all tested initial conditions, and multi-trait eco-evolutionary simulations provide evidence for its evolutionary stability, together supporting a classification as a continuously stable strategy. We close by interpreting this outcome in the context of classical bet-hedging theory and discussing the broader conditions under which biotic and abiotic selective pressures on germination timing are expected to reinforce or oppose one another.

## 2 Ecological Model

We consider a plant population structured into a seed bank of dormant seeds *S* and a class of established adults *A*. Adults reproduce continuously at a density-dependent rate that saturates at carrying capacity *κ*; seeds germinate at per-capita rate *g* and suffer background mortality at rate *d* while dormant. Upon germination, individuals enter the adult stage and are subject to the same background mortality *d*. The germination rate *g* is the focal evolving trait throughout this study.

In the absence of a pathogen, the population dynamics are governed by

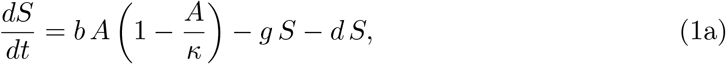

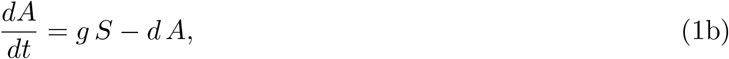

where the logistic seed production term *b A* (1 − *A/κ*) captures intraspecific competition among adults for space and resources. This pathogen-free system establishes a baseline against which the evolutionary consequences of biotic interactions can be assessed.

When a pathogen is present, the adult class is partitioned into susceptible *A* and infected *I* compartments, giving the full three-compartment system

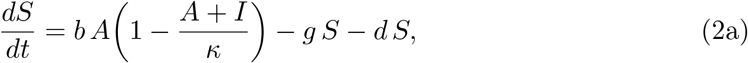

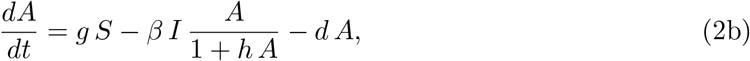

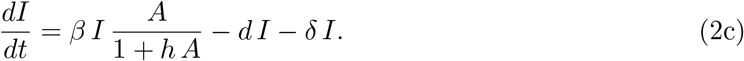

Two features distinguish this system from the pathogen-free model. First, both susceptible and infected adults compete for the same resources, so density-dependent suppression of seed production depends on total adult density, *A* + *I*. Second, pathogen transmission follows a Holling Type II functional response: the per-susceptible rate of new infections saturates at high host density, with handling time *h* capturing the duration of the infection process within a single host. As *h* → 0, the force of infection reduces to the standard mass-action term *β I A*. Infected adults die at the combined rate *d* + *δ*, where *δ* is virulence-associated mortality; no recovery is assumed. Because infection can only be acquired after above-ground establishment, the germination rate *g* governs the rate at which dormant seeds are exposed to pathogen risk upon emergence, and it is precisely this exposure that generates the life-history tension this study is designed to analyse. The model structure is illustrated in Figure 1 and all default parameters values and ranges (chosen to give representative illustrations of the results) are shown in Table 1.

**Table 1.** List of models parameters, default values, and ranges used in the analysis.

| Symbol | Description | Default values | Range |
| --- | --- | --- | --- |
| $b$ | per-capita seed production rate | 0.9 | - |
| $\kappa$ | carrying capacity | 100 | - |
| $g$ | germination rate | 0.9 | $[0, 1]^*$ |
| $d$ | background mortality rate | 0.0001 | - |
| $\beta$ | pathogen transmission rate | 0.3 | $[0, 1]^*$ |
| $h$ | handling time (infection saturation) | 0.15 | - |
| $\delta$ | pathogen virulence rate | 0.8 | $[0, 1]^*$ |

\* Range over which $g$ , $\beta$ , and $\delta$ are examined in this study. Each is a rate with no intrinsic biological upper bound; $[0, 1]$ denotes the interval explored here for tractability, not a physiological/epidemiological ceiling.

**Figure 1.**
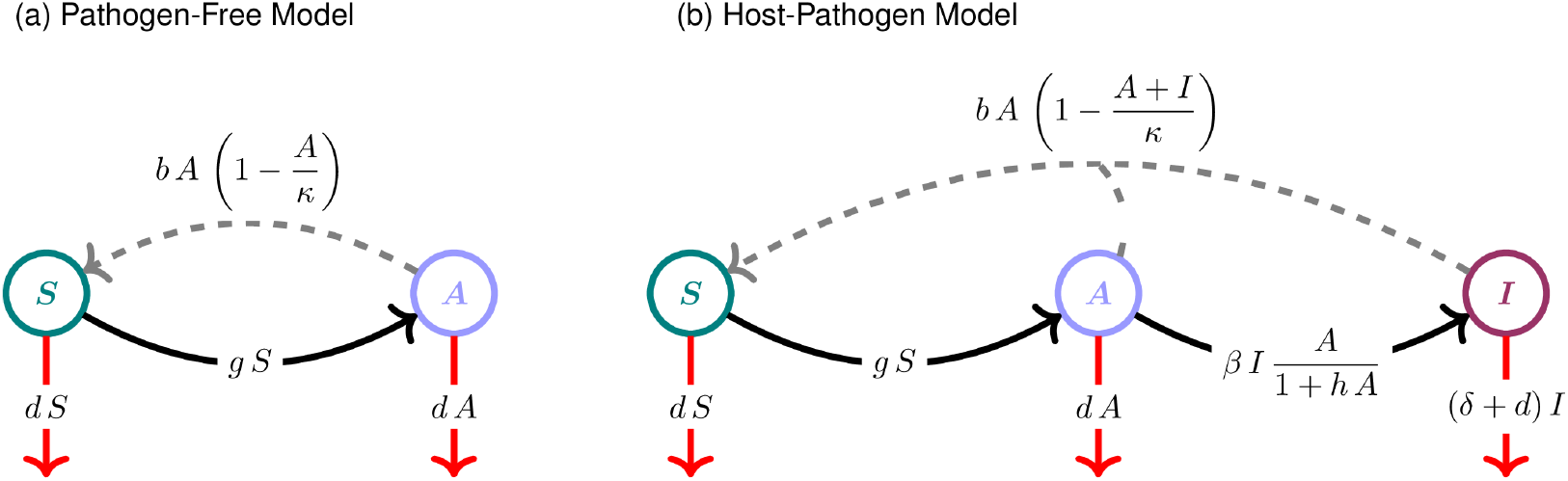
Ecological models schematic illustration. Dashed arrows indicate reproduction; solid arrows indicate transitions between compartments; red arrows indicate mortality.

## 3 Ecological Dynamics

To identify the ecological conditions under which a seed bank can be maintained, we first analyse the equilibria of the pathogen-free model (Equations 1). This system yields two equilibria, summarised in Table 2: a trivial extinction state and a non-trivial coexistence equilibrium. A transcritical bifurcation, a threshold below which the seed cannot persist, occurs at

**Table 2.** List of models equilibria.

|  |  |
| --- | --- |
| **Pathogen-Free Model** |  |
| Equilibrium $(S^*, A^*)$ | |
| Extinction | $(0, 0)$ |
| Coexistence | $\left( \frac{d}{g} \kappa \left( 1 - \frac{d(d+g)}{bg} \right), \kappa \left( 1 - \frac{d(d+g)}{bg} \right) \right)$ |
| **Host-Pathogen Model** |  |
| Equilibrium $(S^*, A^*, I^*)$ | |
| Extinction | $(0, 0, 0)$ |
| Pathogen-Free | $\left( \frac{d}{g} \kappa \left( 1 - \frac{d(d+g)}{bg} \right), \kappa \left( 1 - \frac{d(d+g)}{bg} \right), 0 \right)$ |
| Endemic | $\left( \frac{b(d+\delta)(\kappa(\beta - h(d+\delta)) - \delta)}{(\beta - h(d+\delta))(bg + (d+g)(\beta - h(d+\delta))\kappa)}, \frac{d+\delta}{\beta - h(d+\delta)}, \frac{g((b - b_{\text{Crit}})(\beta - h(d+\delta))\kappa - b(d+\delta))}{(\beta - h(d+\delta))(bg + (d+g)(\beta - h(d+\delta))\kappa)} \right)$ |

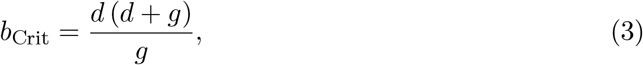

below which seed production is insufficient to offset mortality in both compartments and the population goes extinct; above this threshold the coexistence equilibrium emerges with positive density (Figure 2).

**Figure 2.**
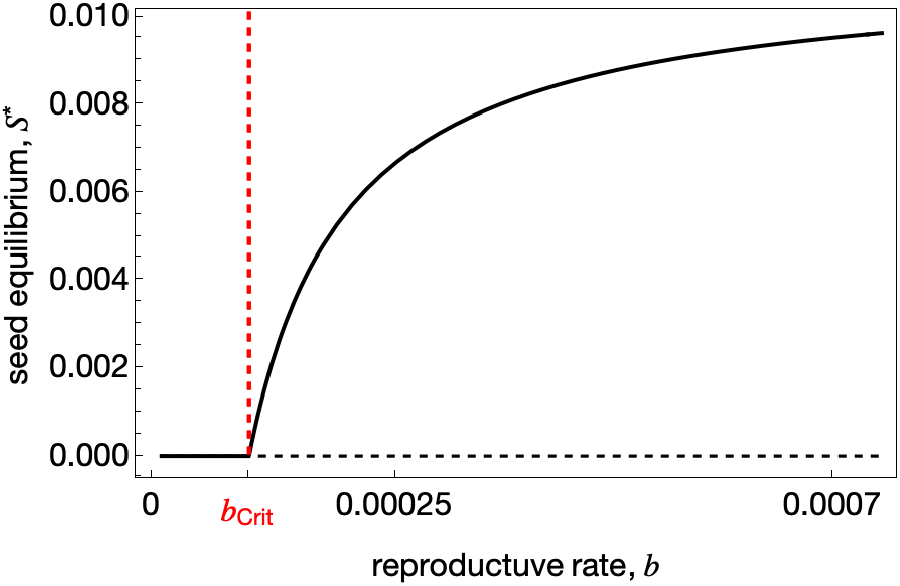
Pathogen-free model bifurcation diagram. Stable and unstable equilibria are shown by solid and dashed curves, respectively. The red dashed vertical line marks the transcritical bifurcation at *b* = *b*_Crit_. Other parameters are the same as Table 1.

The host-pathogen model, Equations (2), admits three biologically distinct equilibria, summarised in Table 2. In addition to the extinction state, there is a pathogen-free equilibrium in which the seed bank and adult class persist while the pathogen is absent, and an endemic equilibrium in which all three compartments coexist. The extinction equilibrium remains unstable for *b > b*_Crit_, as in the pathogen-free model. When transmission is weak, the pathogen fails to invade and the system settles at the pathogen-free equilibrium. The threshold transmission rate at which the pathogen can first invade is given by

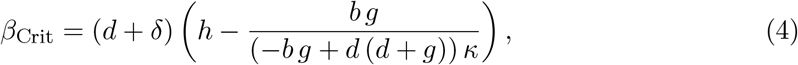

Increasing *β* past *β*_Crit_ destabilises the pathogen-free state via a transcritical bifurcation, giving rise to an endemic equilibrium. The endemic equilibrium is not, however, unconditionally stable: at sufficiently large germination rates it undergoes a Hopf bifurcation, giving rise to sustained periodic dynamics (limit cycles). This generates three qualitatively distinct ecological regimes across parameter space, illustrated in Figure 3a–c: region I, a pathogen-free state; region II, a stable endemic equilibrium; and region III, an oscillatory endemic regime. The boundaries between regions shift systematically across the parameter planes: increasing *β* or decreasing *δ* expands region III at the expense of region II, moving the Hopf boundary to lower values of *g*. The bifurcation diagrams in Figure 3d confirm that this transition is supercritical in both region III cases.

**Figure 3.**
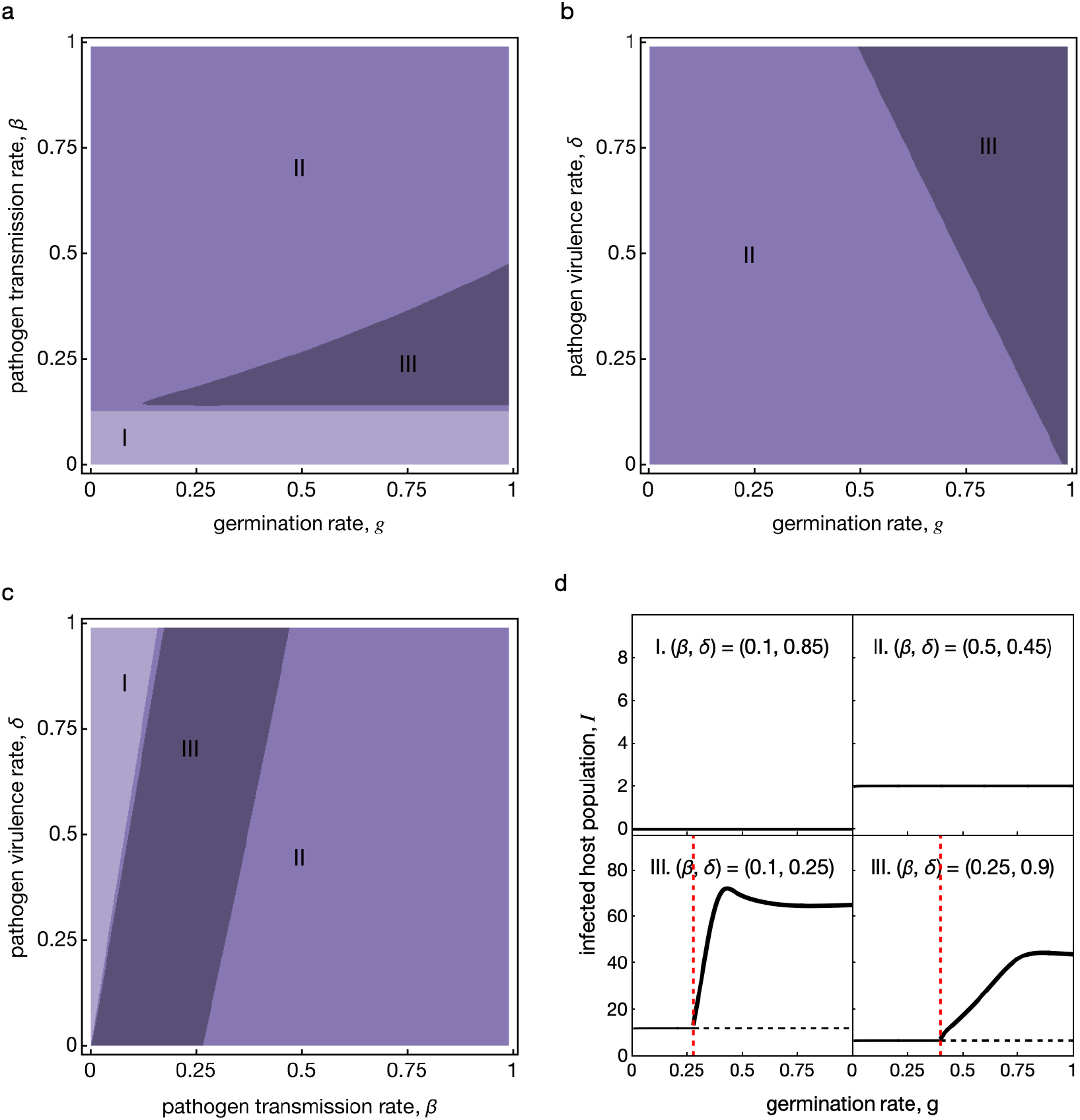
Dynamical regimes of the host-pathogen model and representative bifurcation structure. Panels (a) to (c) show the three qualitatively distinct dynamical regimes — region I (pathogen-free equilibrium), region II (stable endemic equilibrium), and region III (oscillatory endemic) — in the (*g, β*), (*g, δ*), and (*β, δ*) parameter planes, respectively. Panel (d) shows the bifurcation structure in the (*β, δ*) plane: infected host density *I* as a function of germination rate *g* for one representative parameter combination from each region — region I (*β* = 0.1, *δ* = 0.85, top left), region II (*β* = 0.5, *δ* = 0.45, top right), and two region III cases (*β* = 0.1, *δ* = 0.25, bottom left; *β* = 0.25, *δ* = 0.9, bottom right). Solid thin lines indicate the stable equilibriums, solid thick curves show the maximum of *I* along the periodic orbit, and dashed lines show the unstable endemic solution present only in region III. Red dashed vertical lines mark the Hopf bifurcation *g*_Hopf_. Remaining parameters are as in Table 1.

Because *g* simultaneously governs the rate at which dormant seeds are exposed to infection upon germination and determines which dynamical regime the resident population occupies, it acts as both the focal evolving trait and the ecological bifurcation parameter of the system. The dynamical regime the resident population occupies at a given *g* therefore shapes the selection pressure on germination itself, a feedback that is the focus of the following section.

## 4 Evolution of Germination

To determine how natural selection shapes seed germination, we use adaptive dynamics with the invasion analysis detailed in Appendix A. We assume a monomorphic resident population at trait value *g*, and ask whether a rare mutant with trait *g*_*m*_ can grow when introduced at low density into the ecological attractor set by the resident. Because the mutant is rare, it does not alter the resident’s attractor and therefore experiences the resident environment as a fixed background. In regions I & II of Figure 3a–c, where the resident settles to a fixed-point equilibrium, invasion fitness is the leading eigenvalue of the mutant Jacobian evaluated at the resident equilibrium. In region III of Figure 3a–c, the resident attractor is a limit cycle and invasion fitness is the time-averaged per-capita growth rate of the mutant along the resident cycle, the dominant Floquet exponent (see Appendix B). The sign of invasion fitness determines whether the mutant spreads or is excluded; the set of (*g, g*_*m*_) pairs for which invasion is possible is displayed as a pairwise invasibility plots (PIPs).

In the pathogen-free environment (region I of Figure 3a–c), any mutant with *g*_*m*_ *> g* can invade any resident: faster germination translates directly into greater adult recruitment, so the invasion region spans the entire upper triangle of the PIP (see Figure 7a). In the absence of the pathogen, selection on *g* is purely directional: any faster-germinating mutant invades any resident, so there is no interior singular strategy. Within the range *g* ∈ [0, 1] examined here, this directional selection drives the population to the upper edge of the range; absent additional constraints on germination rate (e.g. a physiological ceiling on how quickly a seed can break dormancy), selection would favour ever-increasing *g* without bound. Seeds that delay germination merely postpone reproduction while suffering mortality in the bank, so dormancy confers no fitness benefit when the pathogen is absent. Introducing a stable endemic pathogen (region II of Figure 3a–c) does not alter this outcome: the PIP retains the same upper-triangle structure (see Figure 7d), and selection remains directional with no finite singular strategy, again driving *g* toward the upper edge of the explored range. Although pathogen exposure erodes the adult population, mutants that germinate faster still out-recruit the resident because at a stable endemic equilibrium the benefit of rapid establishment outweighs the elevated infection risk.

The evolutionary outcome changes fundamentally in the limit cycle regime (region III of Figure 3a–c). Because *g* also acts as an ecological bifurcation parameter, the dynamical regime experienced by the mutant depends on where the resident *g* sits relative to the Hopf bifurcation point *g*_Hopf_, creating a qualitative reversal in selection across that threshold. For residents with *g < g*_Hopf_, the upper triangle of the PIP remains shaded: faster-germinating mutants still invade and selection continues to push *g* upward. Once the resident crosses *g*_Hopf_ into region III, the invasion region shifts to the lower triangle so mutants with *g*_*m*_ *< g* now invade, reversing the direction of selection (see Figure 7b & c). The Hopf bifurcation point is therefore a convergent singular strategy: the selection gradient changes sign at *g*^*∗*^ = *g*_Hopf_, driving the population toward this value from either side.

Whether *g*^*∗*^ is also evolutionarily stable (uninvadable) cannot be determined by the standard second-derivative test, because invasion fitness changes character at exactly this point — from an eigenvalue-based criterion (regions I and II) to a Floquet-based criterion (region III). To assess evolutionary stability directly, we initialised a multi-trait ecoevolutionary simulation (see Appendix D) monomorphically at the trait value *g*_*k*_ nearest to *g*_Hopf_ and ran it for *N*_steps_ evolutionary steps. The simulation was repeated under two ecological timescales, *T*_eco_ ∈ {500, 2000}, to verify robustness to the mutation rate (Kim and Ashby, 2026). Evolutionary stability leaves a distinctive signature in the distribution of trait frequencies: a convergence-stable strategy that is also evolutionarily stable produces a narrow, stationary peak concentrated in one or two adjacent traits, whereas a branching point produces a distribution that widens or splits after reaching *g*_Hopf_. In both parameter sets and under both values of *T*_eco_, the final trait distribution remains concentrated in one or two trait classes nearest to *g*_Hopf_ as shown in Figure 5, consistent with mutation-selection balance at a single fitness peak rather than evolutionary branching. We therefore classify *g*_Hopf_ as a continuously stable strategy (CSS). Convergence to this attractor from all initial conditions is further confirmed by numerical trait substitution sequence simulations (Appendix C) and multi-trait trajectories (Appendix D).

**Figure 4.**
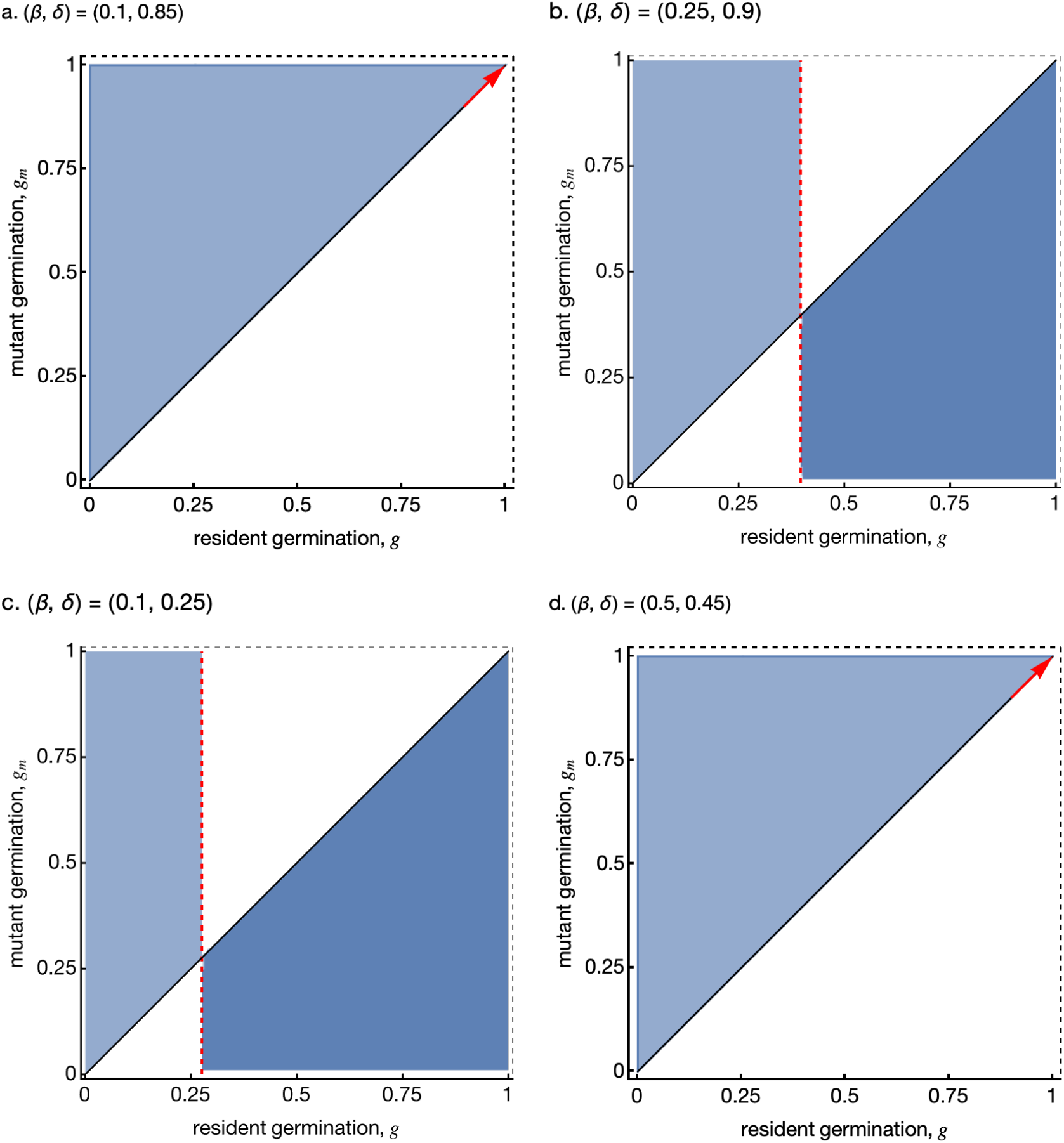
Pairwise invasibility plots (PIPs) for four representative parameter combinations in Figure 3d. In each panel the shaded region indicates where a mutant germination rate *g*_*m*_ can invade a resident *g*. The lighter shaded region corresponds to invasion fitness computed from the leading eigenvalue of the mutant Jacobian at the resident equilibrium, while the darker shaded region is computed from the leading Floquet multiplier of the mutant subsystem along the resident limit cycle. Red dashed vertical lines mark the Hopf bifurcation *g*_Hopf_; the red arrow marks the directional selection (runaway selection). Table 3 provides detailed descriptions of these scenarios. Remaining parameters are the same as in Table 1.

**Figure 5.**
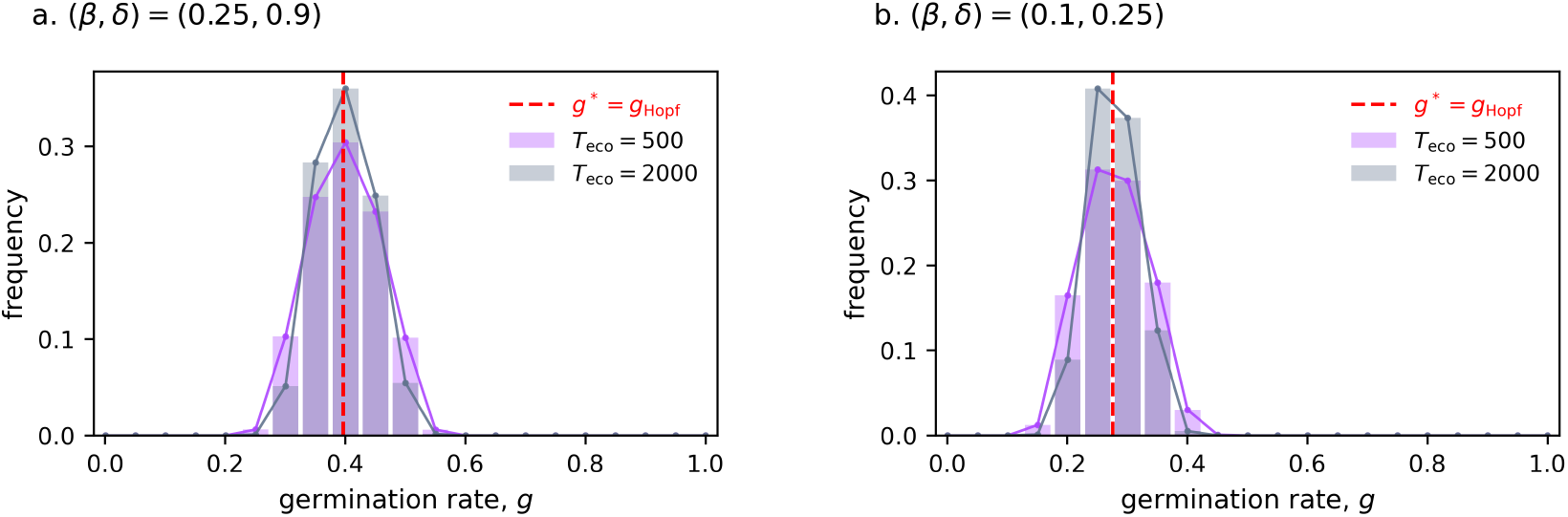
Final trait frequency distributions from multi-trait eco-evolutionary simulations initialised monomorphically at the germination strategy nearest to *g*_Hopf_, under two ecological timescales for two region III parameter sets: (a) (*β, δ*) = (0.25, 0.9) and (b) (*β, δ*) = (0.1, 0.25). Bars show the normalised adult density across germination strategies at the final evolutionary step; the line traces the bar tops.Simulations were run for *T*_evo_ = 750 evolutionary steps with all other parameters as in Table 1.

This reversal is illustrated for two parameter combinations in Figure 7: panel A, *β* = 0.25, *δ* = 0.9, it occurs at *g*_Hopf_ ≈ 0.39: PIP for *g < g*_Hopf_ shows upper-triangle invasion region, while *g > g*_Hopf_ it shows shifted below the diagonal, and in panel C, *β* = 0.1, *δ* = 0.25, the same qualitative reversal is displaced to *g*_Hopf_ ≈ 0.27. In both cases the CSS coincides with the ecological Hopf bifurcation, pinning the population at the boundary between stable and oscillatory dynamics. Table 3 summarizes these outcomes across all three regions.

**Table 3.** Summary of evolutionary scenarios across the three ecological regions. CS = convergence stable; ES = evolutionarily stable; CSS = continuously stable strategy (CS + ES); DIR = directional selection with no finite singular strategy.

|  | Ecological region | Bifurcation | Evol. reversal | Singular strategy | CS | ES | Classification |
| --- | --- | --- | --- | --- | --- | --- | --- |
| I | pathogen-free | none | no | none (unbounded) | N/A | N/A | DIR |
| II | stable endemic | none | no | none (unbounded) | N/A | N/A | DIR |
| III | periodic endemic | supercritical Hopf | yes | interior ($g^* = g_{\text{Hopf}}$) | yes | yes | CSS |

## 5 Discussion

The central result of this study is that endogenously generated pathogen cycles can drive the evolution of seed dormancy to an intermediate, ecologically determined value. When host-pathogen dynamics are limit cycles (region III of Figure 3a–c), selection drives the germination rate toward the Hopf bifurcation boundary (*g*^*∗*^ = *g*_Hopf_). Above this threshold the infection pressure is constant and dormancy confers no advantage: selection becomes purely directional, favouring ever-faster germination with no finite optimum. Within the [0, 1] range explored here this drives the population to the upper edge, *g* = 1; without additional physiological or allocation constraints on germination rate, there is no biologically intrinsic upper bound, and selection would in principle continue to favour still higher rates. Unlike classical dormancy models, in which the selective environment is fixed, here the germination rate shapes its own fitness landscape by controlling whether the resident dynamics are stable or oscillatory.

These results connect naturally to the bet-hedging framework initiated by Cohen (1966). The periodic epidemic peaks are the biotic analogue of abiotic bad years. Cohorts that germinate during prevalence peaks suffer elevated mortality, while those that remain dormant through the peak and emerge during the trough enjoy systematically greater establishment success. Because long-term fitness is governed by the geometric mean growth rate rather than the arithmetic mean, a strategy that accepts reduced average germination in exchange for reduced variance across the cycle is selectively favoured, precisely the bet-hedging logic except driven by biotic rather than abiotic temporal variability. Our Floquet-based invasion criterion captures this directly, since the leading Floquet multiplier represents the geometric-mean growth of the mutant over one resident cycle.

The generality of biotic drivers of life-history evolution extends well beyond delayed germination. Cyclic biotic dynamics are a recurring feature of consumer-resource and host-pathogen systems, and wherever they create periodic variation in juvenile survival they should leave predictable signatures on life-history evolution. The evolution of mast seeding, for instance, may be partly shaped by the build-up and crash of seed and seedling pathogens across high- and low-output years (Kelly, 1994), and the same logic that selects for reduced germination rate in our model could favour episodic, synchronised reproduction over steady annual output (pathogen escape; (Davies and MacPherson, 2024)). More broadly, cyclic pathogen dynamics create the kind of periodic juvenile hazard that life-history theory predicts should shift populations toward slower paces of life, including delayed reproduction, extended dormancy, and along the slow-fast continuum (Promislow and Harvey, 1990; Gaillard et al., 2016). Our results therefore suggest that pathogens, wherever they sustain population cycles, may be an underappreciated driver of pace-of-life evolution across taxa, operating alongside the abiotic drivers that have received most theoretical and empirical attention.

The evolutionary outcome we identify depends on the host-pathogen system producing sustained (limit cycles) rather than damped oscillations, and this requirement is met by a broader range of mechanisms than the Holling Type II transmission assumed here. Distributed latency periods modelled with gamma-distributed delays are well known to generate Hopf bifurcations in otherwise stable endemic models (Blyuss and Kyrychko, 2010), as does waning acquired immunity in SIRS-type systems (Hethcote, 2000); both would be expected to produce qualitatively similar evolutionary dynamics. Cycles arising from predator-prey interactions, competitive boom-bust dynamics, or oscillating mutualists would likewise impose periodic fitness landscapes capable of selecting for dormancy by the same mechanism. The key criterion is that the biotic cycle be sufficiently large in amplitude and long relative to the organism’s generation time to sustain periodic dynamics. The present model is spatially implicit; incorporating spatial structure could reveal how dispersal-dormancy tradeoffs (Vitalis et al., 2013) interact with the temporally driven selection identified here.

## 8 Data availability

No new data were generated or analysed in support of this research. The accompanying *Mathematica* and *Python* code used for analysis is provided with the submission and will be archived on Zenodo upon manuscript acceptance.

## 9 Author contributions statement

M.S. developed the model, conducted all analyses, and drafted the manuscript. A.M. led the planning and supervision of the research and provided critical revisions. J.M.S. contributed to supervision and manuscript editing.

## 10 Acknowledgments

The authors would like to thank Ben Ashby for his helpful feedback on the manuscript. We acknowledge the support of the Natural Sciences and Engineering Research Council of Canada (NSERC).

## Appendix A A Adaptive Dynamics

To determine whether a rare mutant with germination rate *g*_*m*_ can increase when introduced into a resident population at germination rate *g*, we adopt the standard rare-mutant invasion framework of adaptive dynamics (Metz, 1992). The key quantity is the invasion fitness *s*(*g, g*_*m*_), defined as the long-run per-capita growth rate of the mutant when it is sufficiently rare that its dynamical effect on the resident can be neglected. A mutant can invade if and only if *s*(*g, g*_*m*_) *>* 0.

Let (*S*^⋆^, *A*^⋆^, *I*^⋆^) denote the attractor of the resident system at germination rate *g*. When a rare mutant with germination rate *g*_*m*_ is introduced, its classes satisfy

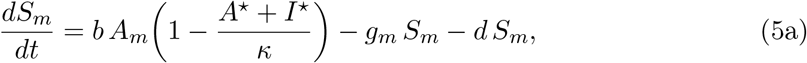

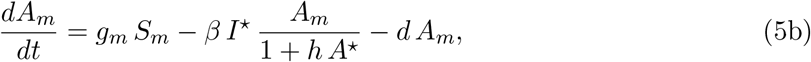

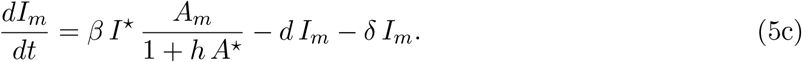

The Jacobian of the mutant (*S*_*m*_, *A*_*m*_, *I*_*m*_) system, evaluated at resident attractor, is

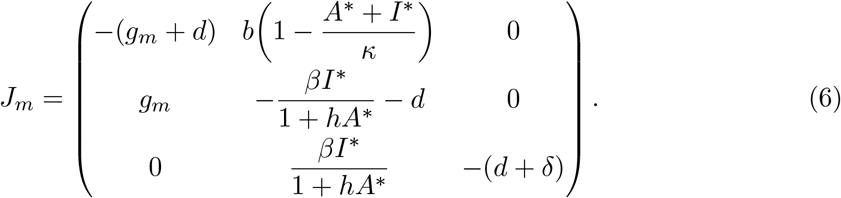

The invasion fitness is the leading eigenvalue of *J*_*m*_, and a rare mutant invades when *s*(*g, g*_*m*_) *>* 0, i.e., when the leading eigenvalue of *J*_*m*_ is positive.

For pathogen-free and endemic equilibrium regions (regions I & II of Figure 3a–c), the resident system converges to a fixed point. And for oscillatory endemic dynamics (region III of Figure 3a–c), the resident system settles on a *T*-periodic limit cycle. The background experienced by the mutant is in this case is limit cycles. In this case, Floquet theory provides the natural framework for computing invasion fitness (see Appendix B).

## B Appendix B Floquet Analysis

Computing invasion fitness in a limit-cycle resident environment requires two numerical ingredients: the period *T* of the resident oscillation, and the monodromy matrix of the mutant subsystem over one cycle.

This approach determines whether small perturbations away from the zero-mutant state grow or decay over one complete resident cycle, and has been widely used in theoretical ecology to characterise selection in periodic environments (Klausmeier, 2008).

First, we determined the period *T* of the resident limit cycle numerically by integrating the resident system past an initial burn-in phase sufficient for transients to decay, then recording the times of successive local peaks of *I*(*t*) in the subsequent observation window; the mean inter-peak interval was taken as *T*, and a confirmed peak time *t*_0_ served as the reference point for the integration interval [*t*_0_, *t*_0_ + *T*].

Next, we constructed the time-dependent Jacobian matrix

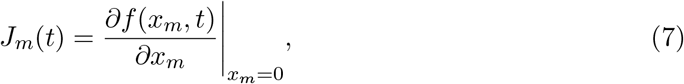

evaluated along the resident limit cycle trajectory (*S*^⋆^, *A*^⋆^, *I*^⋆^), which yields the explicit matrix given in equation (6). We then solved the fundamental matrix equation

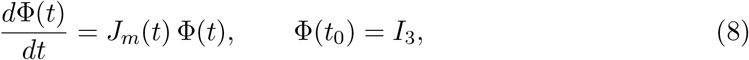

over [*t*_0_, *t*_0_ + *T*] to obtain the monodromy matrix Φ(*t*_0_ + *T*). The leading eigenvalues of Φ(*t*_0_ + *T*), the Floquet multiplier *µ*, determine whether the mutant grows or decays over one cycle.

Finally, we computed the invasion fitness as

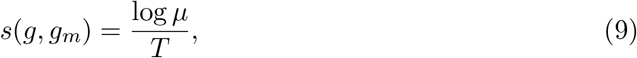

across all (*g, g*_*m*_) pairs in region III of Figure 3a–c. A value *µ >* 1 (equivalently *s >* 0) indicates invasion; *µ <* 1 indicates extinction when rare.

## C Appendix C. Trait Substitution Sequence Simulations

To corroborate the analytical adaptive-dynamics predictions, we implemented a direct numerical simulation of the trait substitution sequence (TSS). Rather than computing invasion fitness analytically, the simulation introduces an explicit rare mutant, integrates the mutant sub-system forward in time on the resident attractor, and accepts the mutant as the new resident if and only if its per-capita population growth rate is positive. Starting from many initial germination rates spanning the full parameter space, every trajectory converged to the Hopf bifurcation threshold *g*^*∗*^ = *g*_Hopf_ (Figure 6), confirming that *g*_Hopf_ is the evolutionary attractor predicted by the theory.

**Figure 6.**
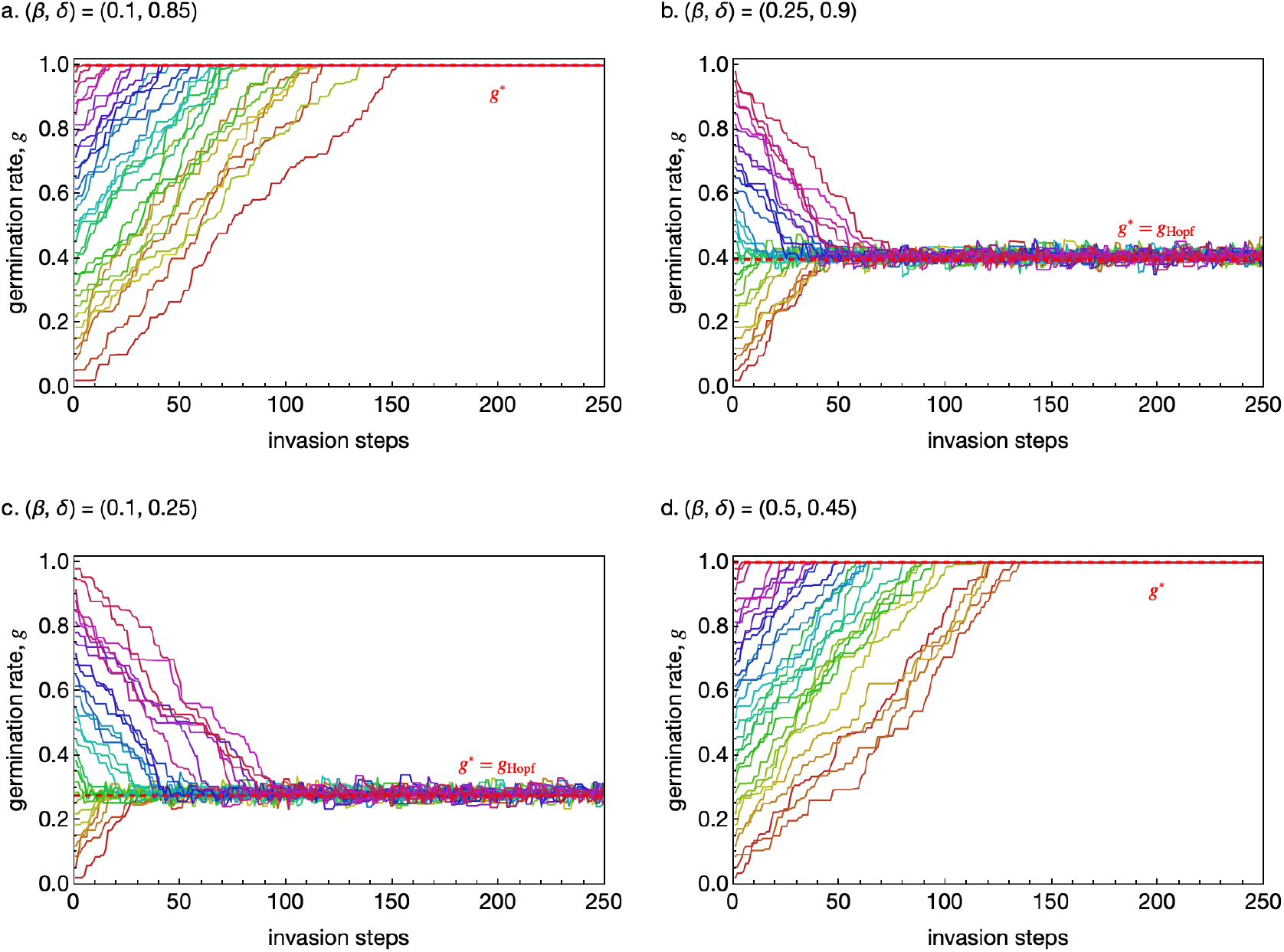
Trait substitution sequence (TSS) simulations corresponding to the parameter combinations in Figure 7. Each coloured line tracks the resident germination rate *g* across 250 substitution steps; the red dashed line marks the convergence-stable singular strategy. Panels a–d use the same parameter sets as the respective PIP panels. In panels a and d (regions I and II) all trajectories converge to *g*^*∗*^ = 1, the upper edge of the trait range [0, 1] explored here — reflecting unbounded directional selection rather than a true convergencestable strategy — while in panels b and c (region III) trajectories initiated both above and below *g*_Hopf_ converge to *g*^*∗*^ = *g*_Hopf_, confirming the analytical prediction.

The TSS simulation is built from four components: a resident integrator, a mutant invasion test, a single substitution step, and a trajectory runner. The logic of each step is described below and summarised in Algorithm 1.

For a given resident germination rate *g*, system (2) is integrated numerically past a burn-in phase sufficient for transients to decay, after which the attractor is classified as either a stable equilibrium (regions I and II) or a limit cycle (region III), along with the relevant diagnostics needed by the invasion test.

Given a mutant germination rate *g*_*m*_ and the resident attractor association, the mutant sub-system is integrated over a test window with the resident trajectory fixed as a time-varying background (linearised-in-mutant equations; density-dependent terms in *S*_*m*_, *A*_*m*_, *I*_*m*_). The initial mutant densities are set to a small fraction *ε* = 0.02 of the resident state at the start of the test window. The test window spans 5 time units (equilibrium case) or 5 *T* (limit-cycle case). The per-capita growth rate of the total mutant population is computed as

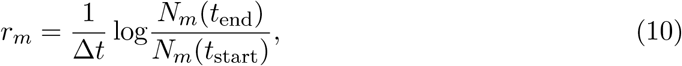

where *N*_*m*_ = *S*_*m*_ + *A*_*m*_ + *I*_*m*_ and the mutant is declared successful if *r*_*m*_ *>* 0.

At each step, a candidate mutant trait is drawn from a normal distribution centred on the current resident: *g*_*m*_ ~*g* + *N*(0, *σ*^2^) with *σ* = 0.02, clipped to the interval [0.001, 1.0]. The upper clip at *g*_*m*_ = 1 mirrors the [0, 1] range used throughout the analytical treatment; because selection on *g* is directional and unbounded in regions I and II, without this clip the resident trait would increase indefinitely rather than converging to a finite value. If the invasion test returns success, *g*_*m*_ replaces *g*; otherwise the resident trait is unchanged. Starting from an initial trait value *g*_0_, the simulation iterates the substitution step for *N*_steps_ = 250 steps, caching the resident integration between consecutive steps at which the resident trait is unchanged, so that the expensive ODE solve is not repeated for failed invasion attempts. The function returns the complete history (*g*_0_, *g*_1_, …, *g*_250_).

We ran 30 independent trajectories with initial germination rates drawn uniformly from the interval [0.02, 0.98]. For each of two (*β, δ*)-parameter sets, all trajectories converged to *g*^*∗*^ = *g*_Hopf_ within the 250-step observation window (Figure 6). Trajectories initiated in region II (*g*_0_ *> g*_Hopf_, stable endemic equilibrium) drifted monotonically downward toward *g*_Hopf_, while those initiated in region III (*g*_0_ *< g*_Hopf_, limit cycle) drifted upward, consistent with the convergence-stability analysis.

## D Appendix D. Multi-Trait Eco-Evolutionary Simulations

To assess evolutionary stability at the singular strategy *g*^*∗*^ = *g*_Hopf_, where the standard second-derivative test is inapplicable due to the degeneracy of the invasion fitness function at the Hopf boundary, we implemented a multi-trait eco-evolutionary simulation following the design of Kim and Ashby (2026).

Rather than tracking a single monomorphic resident, this simulation maintains all *m* germination strategies simultaneously in a shared ecology. Mutations transfer density between adjacent strategies, and selection acts through the ecological dynamics without any explicit computation of invasion fitness.

The trait space [0.001, 1.0] is discretised into *m* = 21 equally spaced germination rates *g*_1_ *< g*_2_ *<* · · · *< g*_*m*_, with trait step size Δ*g* ≈ 0.05. Each strategy *k* ∈ {1, 2, · · ·, *m*} maintains its own compartments (*S*_*k*_, *A*_*k*_, *I*_*k*_), giving a 3*m*-dimensional state vector. All strategies share the same pathogen pool and carrying capacity, so they compete ecologically through the shared quantity ∑_*k*_ *A*_*k*_ + ∑_*k*_ *I*_*k*_. The governing equations for strategy *k* are

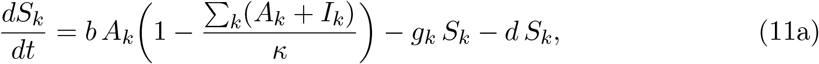

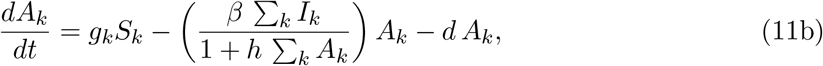

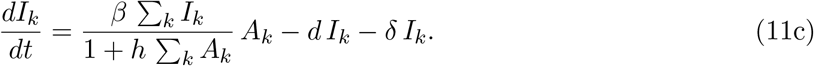

### Algorithm 1

Trait substitution sequence simulation.

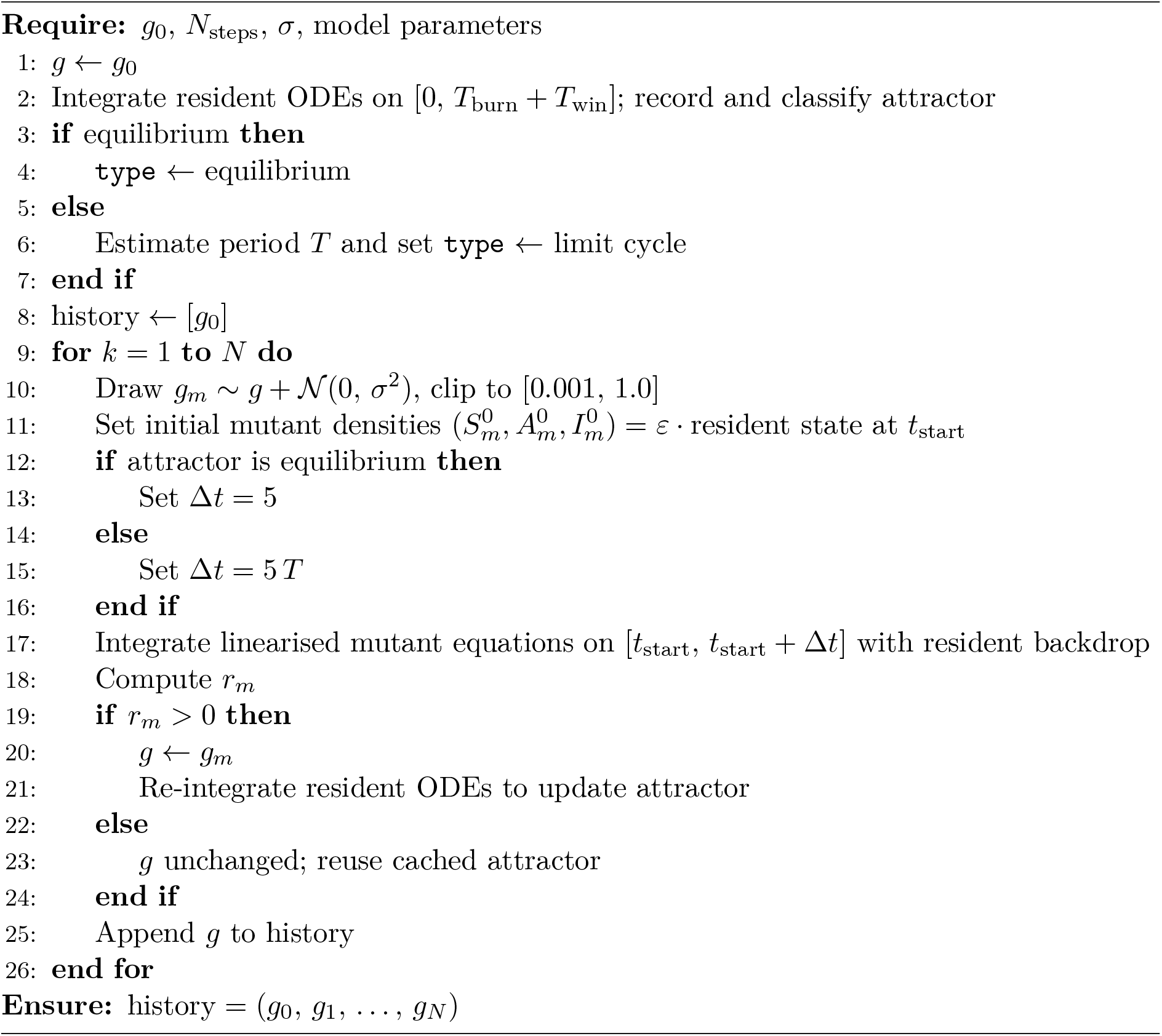

The system is integrated using a fixed-step fourth-order Runge-Kutta scheme with step size Δ*t* = 0.5. Any compartment whose density falls below the extinction threshold *ε* = 10^−9^ is set to zero.

At each evolutionary step, a single mutation event is introduced following Kim and Ashby (2026). A progenitor strategy *k*_*p*_ is selected with probability proportional to its current adult density 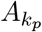, so that reproductively active strategies generate mutations more frequently (analogous to reproducing stage in Kim and Ashby (2026)’s model). The mutant strategy is one discrete step away from the progenitor, *k*_*m*_ = *k*_*p*_ *±* 1 with equal probability, reflecting at the boundaries *g*_1_ and *g*_*m*_. A fixed fraction *η* = 0.01 of the progenitor’s seed-bank density 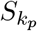 is transferred to 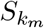. The adult and infected compartments of the mutant 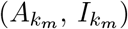 start at zero and build up through subsequent ecological dynamics, analogous to Kim and Ashby (2026), where only the free parasite is transferred and the infected class develops naturally. The process repeats for *N*_step_ = 750 evolutionary steps.

For each of the two region III parameter sets (Figure b–c), 20 independent trajectories were initialised monomorphically at germination rates *g*_0_ evenly distributed across [0.001, 1.0], so that trajectories began both above and below *g*_Hopf_. The weighted mean germination rate 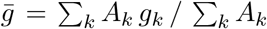 was recorded at each step. Trajectories initiated above and below *g*_Hopf_ both converged to 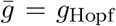 as shown in Figure 7, confirming the convergence-stability result from the analytical invasion analysis.

**Figure 7.**
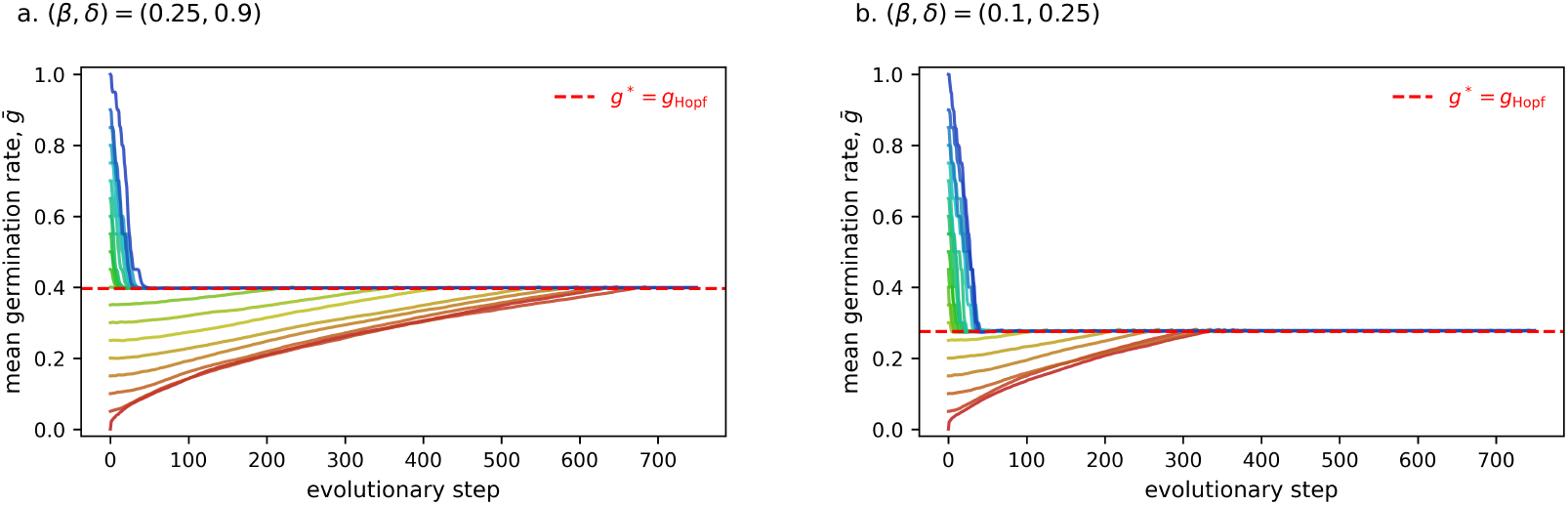
Eco-evolutionary simulations of germination rate for two region III parameter sets (Figure b–c). Each coloured line tracks the weighted mean germination rate 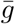 across 750 evolutionary steps, starting from a different initial germination rate. Trajectories initiated both above and below *g*_Hopf_ converge to *g*^*∗*^ = *g*_Hopf_ (red dashed line), confirming convergence stability.

## References

A. Best and B. Ashby. How do fluctuating ecological dynamics impact the evolution of hosts and parasites? Philosophical Transactions of the Royal Society B: Biological Sciences, 378(1873):20220006, 2023. doi: 10.1098/rstb.2022.0006.

Konstantin B. Blyuss and Yuliya N. Kyrychko. Stability and bifurcations in an epidemic model with varying immunity period. Bulletin of Mathematical Biology, 72(2):490–505, 2010. doi: 10.1007/s11538-009-9458-y.

Dan Cohen. Optimizing reproduction in a randomly varying environment. Journal of Theoretical Biology, 12(1):119–129, 1966. doi: 10.1016/0022-5193(66)90188-3.

T. Jonathan Davies and Ailene MacPherson. Seed masting as a mechanism for escape from pathogens. Current Biology, 34(4):R120–R125, 2024. doi: 10.1016/j.cub.2023.12.027.

William E. Finch-Savage and Gerhard Leubner-Metzger. Seed dormancy and the control of germination. New Phytologist, 171(3):501–523, 2006. doi: 10.1111/j.1469-8137.2006.01787.x.

J.-M. Gaillard, J.-F. Lemaître, V. Berger, C. Bonenfant, S. Devillard, M. Douhard, M. Gamelon, F. Plard, and J.-D. Lebreton. Life Histories, Axes of Variation in. In Encyclopedia of Evolutionary Biology, pages 312–323. Elsevier, 2016. doi: 10.1016/B978-0-12-800049-6.00085-8.

Jennifer R. Gremer and D. Lawrence Venable. Bet hedging in desert winter annual plants: optimal germination strategies in a variable environment. Ecology Letters, 17(3):380–387, 2014. doi: 10.1111/ele.12241.

Herbert W. Hethcote. The Mathematics of Infectious Diseases. SIAM Review, 42(4):599–653, 2000. doi: 10.1137/S0036144500371907.

C. S. Holling. The components of predation as revealed by a study of small-mammal predation of the european pine sawfly. The Canadian Entomologist, 91(5):293–320, 1959. doi: 10.4039/Ent91293-5.

Dave Kelly. The evolutionary ecology of mast seeding. Trends in Ecology & Evolution, 9 (12):465–470, 1994. doi: 10.1016/0169-5347(94)90310-7.

Yoon Soo Kim and Ben Ashby. Coevolutionary cycling in allele frequencies and the evolution of virulence. Evolution, 80(1):282–290, 2026. doi: 10.1093/evolut/qpaf224.

Christopher A. Klausmeier. Floquet theory: a useful tool for understanding nonequilibrium dynamics. Theoretical Ecology, 1(3):153–161, 2008. doi: 10.1007/s12080-008-0016-2.

J A J Metz. How Should We Define’Fitness’for General Ecological Scenarios? TREE, 7(6), 1992.

D. E. L. Promislow and P. H. Harvey. Living fast and dying young: A comparative analysis of life-history variation among mammals. Journal of Zoology, 220(3):417–437, 1990. doi: 10.1111/j.1469-7998.1990.tb04316.x.

Mark Rees. Delayed Germination of Seeds: A Look at the Effects of Adult Longevity, the Timing of Reproduction, and Population Age/Stage Structure. The American Naturalist, 144(1):43–64, 1994. doi: 10.1086/285660.

M. F. Scott and S. P. Otto. Why wait? Three mechanisms selecting for environment-dependent developmental delays. Journal of Evolutionary Biology, 27(10):2219–2232, 2014. doi: 10.1111/jeb.12474.

Shripad Tuljapurkar and Pamela Wiener. Escape in time: stay young or age gracefully? Ecological Modelling, 133(1–2):143–159, 2000. doi: 10.1016/S0304-3800(00)00288-X.

D. Lawrence Venable and Joel S. Brown. The Selective Interactions of Dispersal, Dormancy, and Seed Size as Adaptations for Reducing Risk in Variable Environments. The American Naturalist, 131(3):360–384, 1988. doi: 10.1086/284795.

Renaud Vitalis, François Rousset, Yutaka Kobayashi, Isabelle Olivieri, and Sylvain Gandon. The joint evolution of dispersal and dormancy in a metapopulation with local extinctions and kin competition. Evolution, 67(6):1676–1691, 2013. doi: 10.1111/evo.12069.

